# *Cgas* deficiency promotes tumor growth by supporting B cell persistence and angiogenesis

**DOI:** 10.1101/2024.09.24.614699

**Authors:** Papasara Chantawichitwong, Sarinya Kumpunya, Tossapon Wongtangprasert, Peerapat Visitchanakun, Trairak Pisitkun, Prapaporn Pisitkun

## Abstract

The cGAS sensor activates STING/IFN signaling, which is crucial for immune defense against pathogens and triggers inflammation in autoimmune diseases and antitumor responses. This study investigated the cGAS-mediated immune response in tumorigenesis using the MC-38 tumor model. *Cgas^-/-^* mice exhibited significantly larger tumors and lower survival rates than wild-type (WT) mice. Tumors in *Cgas^-/-^* mice showed increased fibrosis and neovascularity. WT mice mounted a more robust T-cell-mediated antitumor response, with higher levels of NK and effector T cells, while *Cgas^-/-^* mice showed an expansion of B cells, including regulatory B cells producing IL-10. B cells from tumor-bearing *Cgas^-/-^* mice survived better in the tumor- conditioned medium than those from WT mice. B cell depletion significantly reduced tumor size in WT mice but had minimal effect in *Cgas^-/-^* mice, where fibrosis and tumor vasculature persisted. Despite B cell depletion, B cells remained in the tumors of *Cgas^-/-^* mice, in contrast to WT mice, where their reduction correlated with an increase in CD8^+^ infiltrating cells. Expression of *Tlr7* and *Tlr9* remained elevated and unaffected by B cell depletion in *Cgas^-/-^* tumors, while *Baff* expression was higher and further increased after B cell depletion. *Cgas^-/-^* B cells promoted angiogenesis, as indicated by enhanced endothelial tube formation. In summary, cGAS deficiency fosters a tumor microenvironment that supports B cell survival, promotes a pro-tumor immune environment, and enhances angiogenesis, contributing to tumor progression.

## Introduction

The tumor microenvironment (TME) is a crucial factor that influences cancer progression and is composed of extracellular matrix (ECM), proliferating tumor cells, stromal cells, blood vessels, infiltrated inflammatory immune cells, and other associated tissue cells (1). The type of immune cell infiltration is associated with the prognosis of cancer (2). High levels of infiltrating T cells are associated with a good prognosis in many types of cancer (3, 4). However, the role of B cells in the pathogenesis of solid tumors is diverse between cancer types (5).

Cyclic GMP-AMP synthase (cGAS) is a cytosolic pattern recognition receptor that detects intracellular pathogens and results in the conversion of ATP and GTP into 2,3-cyclic GMP-AMP (cGAMP) to activate the endoplasmic reticulum (ER)-resident adaptor protein STING and induce interferon (IFN) production(6). The cGAS/STING pathway is an inflammatory driver resulting from genomic instability and DNA damage caused by carcinogens or radiation therapy (7, 8). Nucleic acids derived from cancer cells can activate the cGAS/STING pathway and lead to antitumor immunity (7, 9). cGAS/STING signaling is activated by cytosolic DNA in tumor cells or cGAMP in neighboring cells (10).

The persistence of inflammation promotes tumor progression (11). Cancer cells and the microbiomes in tumors can modulate the microenvironment to create an inflammation that suits cancer development (12). Chronic inflammation is associated with tissue fibrosis (13, 14). TGF-β could be a proinflammatory cytokine under chronic injury conditions (14, 15). TGF-β activation could also induce EMT, which could cause malignant tumor growth (15). In addition, TGF-β can promote angiogenesis through SMAD transcriptional regulation (16). TGF-β ligands are secreted by almost all cell types, including epithelial cells, fibroblasts, and immune cells, and they are inactive and stored in the TME (17).

Activation of the cGAS/STING pathway increases the strength of dendritic cells (DCs), presenting tumor antigens to CD8^+^ T cells to create an antitumor effect (18, 19). However, inhibition of STING signaling in B cells reduces tumor burden in multiple orthotopic tumor cell lines (20). Autoimmune features and rheumatic manifestations frequently occur in cancer patients (21). Anti-oncoprotein and antitumor suppression gene antigens, detected as autoantibodies, are seen before cancer diagnosis or in the early stages of malignant disease (21). In addition to cancer, STING signaling also initiates autoimmune lupus disease (22). However, without cGAS, dsDNA can activate the cGAS/inflammasome pathway and cause more inflammation in pristane-induced lupus (23).

B cells influence both pro- and antitumor immunity. Studies show a favorable outcome with tumor-infiltrating B cells in cancer (24, 25). However, not all B cell subsets promote antitumor responses; in lung cancer, Bregs can suppress it (26). B cells can also drive tumor growth through immunoglobulin production and chronic inflammation via FcγR activation (27). Research indicates *Cgas*-deficient mice are prone to develop autoimmunity in pristane-induced lupus, suggesting they develop chronic inflammation and hyperreactive B cells upon stimulation (23, 28).

In the MC-38 cancer model using C57BL/6 mice, enhanced tumor progression was observed in *Cgas^-/-^* mice, accompanied by an increase in IL10^+^ B cells, higher survival of B cells with tumor antigens, and antinuclear antibody, indicating an autoreactive B cell response. Depleting B cells reduced tumor volume in wild-type (WT) but not in *Cgas^-/-^* mice, with fibrosis scores higher in the latter. Additionally, B cells from *Cgas^-/-^* mice were more resistant to B cell depletion and more effective in promoting endothelial cell tube formation than those from WT mice. This data suggests that B regulatory cells are crucial in tumor progression, with CGAS signaling inhibition leading to a proinflammatory tumor microenvironment and accelerated tumor growth. *Cgas^-/-^* B cells enhance angiogenesis, and B cell depletion could serve as an additional immunotherapy approach in certain cancers. Identifying patients with specific biomarkers, such as ANA, could help select candidates for treatment, highlighting the need for further research.

## Materials and Methods

### Mouse model

The cGAS-deficient C57BL/6 mice were provided by Professor Søren Riis Paludan (Aarhus University, Aarhus, Denmark), and the WT mice were purchased from Nomura Siam International, Bangkok, Thailand. MC-38 cell lines were cultured in DMEM with 10% FBS, 2 mM L-glutamine, 0.1 mM nonessential amino acids, 1 mM sodium pyruvate, 10 mM Hepes, and 50 µg/ml gentamicin sulfate 1x Antibiotic-Antimycotic (Gibco-Thermo Fisher Scientific, MA, USA) at 37°C and 5% CO₂. Age-matched 6–8-week-old mice were inoculated subcutaneously (right hind flank) with 3x10^5^ MC-38 cells in 100 µl PBS. The mice were observed for 30-45 days to analyze tumor volume and survival. Tumor volume was recorded by digital vernier caliper measurement and calculated using the following formula: tumor volume = (width^2^ x length)/2. The analysis was carried out blindly by two investigators to reduce bias. The animal protocols were approved by the Institutional Animal Care and Use Committees (IACUC) of the Faculty of Medicine, Chulalongkorn University (018/2563). All methods were performed following the ARRIVE guidelines. All procedures were performed following the relevant guidelines and regulations.

### Histological analysis

The tumor was harvested and preserved in formalin-fixed paraffin-embedded (FFPE) tissue. The 5 μm sections of the tumor were stained with hematoxylin and eosin (H&E) to observe cellular structure. Fibrosis score was evaluated by the following criteria, Grade 0 = no fibrosis; mainly tumor cells in the area, Grade 1 = low fibrosis; with an increased fibroblast surrounding carcinoma, Grade 2 = medium fibrosis; an intermediate between grade 1 and grade 3, Grade 3 = high fibrosis; mainly comprised of fibers with few tumor cells in the area. The tumor sections were stained with anti-Vimentin (dilution 1:500; ab92547, Abcam), anti-CD31 (dilution 1:200; ab182981, Abcam), anti-α-SMA (dilution 1:200; #19245, Cell signaling), anti- VEGFR2 (dilution 1:200; #2479, Cell signaling), anti-Ki-67 (dilution 1:400; D3B5; #9129, Cell Signaling) and DAPI for immunofluorescence analysis then visualized under a ZEISS LSM 800. The mean fluorescent intensity was evaluated in 5 fields for each slide by image J and the Ki-67 index was calculated by (positive Ki-67 cells/total positive nuclear staining cells) x 100.

### Flow cytometry analysis

Splenocytes (1 x 10^6^ cells) were resuspended in staining buffer (0.5% BSA in PBS and 0.09% azide) and stained with fluorophore-conjugated antibodies, including anti-CD3 (145-2C11; #100312), CD4 (GK1.5; #100423), CD8 (53-6.7; #100708), CD44 (IM7; #103032), CD62L (MEL-14; #104418), CD56 (NCAM; #318304), B220 (RA3-6B2; #103222), F4/80 (BM8; #123110), CD11b (M1/70; #101230), CD11c (N418; #114310), CD21 (Bu32; #354908), CD23 (B3B4; #101614), CD19 (6D5; #115512), Ly6c (HK1.4; #128006), and Ly6g (1A8; #127608) (Biolegend, San Diego, CA, USA). For intracellular staining, splenocytes were incubated with 25 ng/ml PMA, 1 μg/ml ionomycin (Sigma-Aldrich, Darmstadt, Germany), and 1x GolgiPlug (Brefeldin A, Biolegend, San Diego, CA, USA) for 4 hours at 37 °C and 5% CO₂. Cell surface antigen staining was performed. Cells were fixed in 200 μl of fixation buffer (Biolegend, San Diego, CA, USA) in the dark at 4 °C overnight. Next, the cells were washed 3 times, resuspended in 1x permeabilization buffer (Biolegend, San Diego, CA, USA), and stained with anti-IL-10 (JES5-16E3; #505026) and IFN-γ (XMG1.2; #505821). Flow cytometry was performed using a BD® LSR II Flow Cytometer (BD Biosciences) and analysis by FlowJo software version 10.9.0 (BD, Ashland, OR, USA).

### Measurement of an antinuclear antibody (ANA) and immunoglobulin

ANA from sera was detected using HEp2 substrate slides (coated with HEp-20- 10 cells) at a 1:50 dilution (EUROIMMUN AG, Luebeck, Germany). Fluorescence signaling was visualized by a ZEISS LSM 800 (Carl Zeiss, Germany). The plates were coated overnight at 4 °C with 20 ng/well mouse anti-IgG. After incubation, the plates were washed 3 times with PBST (0.1% Tween (v/v) in 1x PBS). Serum samples (1:60000) in 10% BSA/TBST were placed on the plate and then incubated at 37 °C for 1 hour. Goat anti-mouse IgG conjugated to HRP (1: 8000) was added and set at 37 °C for 1 hour. TMB substrate was added and incubated at room temperature for 20 mins. The reaction was stopped, and the intensity was measured at 450 nm.

### Carboxyfluorescein succinimidyl ester (CFSE) labeling

The isolated B cells (1 x 10^6^ cells) were incubated in prewarmed (37 °C) 1x PBS with 0.5 μM CFSE (Biolegend, San Diego, CA, USA) at 37 °C in a 5% CO₂ incubator for 10 minutes. Next, the reactions of the labeling cells were quenched with 10 volumes of ice-cold complete medium and centrifuged for 5 minutes at 1,500 rpm and 4°C. The CFSE-labeled cells were centrifuged and washed in a complete RPMI-1640 medium. The cells were detected for proliferation by flow cytometry.

### B cell isolation for *in vitro* cell culture

Negative selection isolated naive B cells from mouse spleens using the EasySep™ Mouse B Cell Isolation Kit (cat. 19854, STEMCELL Technologies). The isolated B cells (1 x 10^5^) were cultured in complete RPMI-1640 medium (supplemented with 10% FBS, 1 mM sodium pyruvate, 2 mM L-glutamine, 10 mM HEPES, 0.1 mM nonessential amino acids, 1x Antibiotic-Antimycotic (Gibco) and 50 μM 2-mercaptoethanol (Sigma-Aldrich)) on a 96-well plate in 5% CO_2_ at 37°C for 48 hours. 1 x 10^6^ MC-38 cells were cultured in a T75 flask in 10 ml of complete RPMI- 1640 medium for 48 hours at 37 °C and 5% CO_2_. The supernatant was collected and centrifuged at 1500×g for 15 mins at RT and stored at -80°C before being used as tumor-conditioned media (TCM).

### B cell stimulation *in vitro*

The B cells (1 x 10^5^ cells) were cultured in complete RPMI-1640 medium and were transfected with 1 mg/ml cGAS agonist (G3-ended Y-form Short DNA) (InvivoGen, San Diego, USA) and LyoVec™ (InvivoGen, San Diego, USA). TCM (1:2 dilution with complete RPMI-1640) and complete RPMI-1640 medium were used as the control for the TCM experiment. After 48 hours of incubation, the cells were stained for anti-CD19 (Biolegend) and Live/Dead with Fixable Viability Dye eFluor™ 780 (#65-0865-14, Thermo Fisher Scientific, MA, USA) and then examined by flow cytometry.

### B cell depletion *in vivo*

C57BL/6 mice (6-8 weeks old) were once intravenously injected with 0.25 mg of Ultra-LEAF™ purified anti-mouse CD-20 antibody (SA271G2; #152115, BioLegend, San Diego, CA, USA) 3 days before MC-38 subcutaneous tumor inoculation. At day 0 and day 7, mouse blood was collected, and the cells were stained with anti-CD45 (30- F11; #103114), anti-CD19 (Biolegend), and Live/Dead with Fixable Viability Dye eFluor™ 780 (Thermo Fisher Scientific) to analyze the percentage of B cells by flow cytometry.

### Multiplex immunofluorescence staining

Per the manufacturer’s instructions, staining was performed with the Opal Kit for multiplex IHC (Akoya Bioscience). Briefly, the FFPE tissue section was deparaffinized, and rehydration and blocking endogenous peroxidase were performed. Sections were washed with 1x TBST and blocked with blocking/antibody diluent for 10 minutes before being incubated with primary rabbit monoclonal antibodies, including anti-CD8 (D4W2Z; #98941, Cell Signaling) and anti-CD19 (EPR23174-145; # ab245235, Abcam). Next, the sections were incubated with Opal Polymer HRP Ms + Rb for 10 minutes, followed by an Opal fluorophore (including Opal 540 and Opal 570) for 10 minutes. Each antibody staining was paired with the Opal fluorophore.

Finally, the tissues were stained with DAPI (4′,6-diamidino-2-phenylindole) for 5 minutes and mounted in ProLong™ Diamond Antifade Mountant (Thermo Fisher Scientific, MA, USA).

### Spectral library and imaging

A spectral library was created for multispectral image analysis visualization and fluorophore extraction. A spleen from the WT mouse was used as a control and was stained using the same antibodies stained in samples for an expression marker linked to each Opal fluorophore tyramide. Single-color staining was performed along with the samples but without DAPI to obtain an abundant signal with each fluorophore. The Vectra Polaris Automated Quantitative Pathology Imaging System (Akoya Biosciences) was used for multispectral imaging at 20× magnification. Then, all the slide images were loaded into inForm® Tissue Analysis Software (Version 2.6.0, Akoya Bioscience) to perform batch processing and image analysis for cell density profiles. The various markers were characterized and quantified using the same algorithm.

### Quantitative real-time PCR

Total RNA was extracted from the spleens and tumors using an RNeasy RNA extraction kit (Qiagen, CA) following the manufacturer’s instructions. Total RNA was treated with DNase I (QIagen, CA) to eliminate contaminating genomic DNA. The RNA was then converted into cDNA using an iscriptTM cDNA synthesis kit (Bio-rad). All cDNA samples were stored at -80 ^0^C. Total RNA was reverse transcribed using iScript™ Reverse Transcription supermix (Bio-rad). qRT-PCR was performed using 50 ng of cDNA and SYBR green dye on QuantStudio™ 5 Real-Time PCR System (Applied Biosystems). Relative quantities (Δct values) were obtained by normalizing against the *actin* of the spleen control sample. The primer sequences are shown in S1 Table.

### Tube formation assay

Cultrex RGF BME type R1 was coated in 96-well plates at 37 °C for 1 h. Then, 15,000 HUVECs were seeded in the EBM-2 medium with 2% FBS on Matrigel. The tube formation ability of HUVECs was observed at 4h, 8h, and 24h under microscopy. Image J software quantified the number of tubes and nodes of the tubular structures.

EGF (10 ng/mL) and VEGF (10 ng/mL) were added to the culture medium as a positive control. The HUVECs were cultured with TCM (1:1) and murine’s splenic-B cell (1 x 10^5^ cells) to evaluate the effect of *Cgas^-/-^* B cell on the ability of endothelial cells to form a tubular structure.

### Statistical analysis

Statistical tests were performed with GraphPad Prism 8.0 (GraphPad Software).

The unpaired two-tailed Student’s t-test was used to compare two experimental groups. Survival data were analyzed with a log-rank (Mantel-Cox) test. Tumor volume records were analyzed with two-way ANOVA. Data are shown as the mean ± SEM, and P values < 0.05 were considered statistically significant. The sample size was calculated with a 95% confidence interval.

## Results

### The absence of cGAS promotes progressive tumor formation and poor survival

The *Cgas*^-/-^ mice significantly increased tumor growth more than WT mice after 28 days of MC-38 tumor inoculation (Fig 1A), whereas WT mice showed a significantly higher rate of survival compared to *Cgas*^-/-^ mice after tumor injection at day 39 (*p < 0.0001*) (Fig 1B). *Cgas*^-/-^ mice tumors showed higher neovascularity and fibrosis than WT mice (Figs 1C and 1D). The histopathology revealed a higher fibrosis score in *Cgas*^-/-^ mice (Fig 1E). Then, we analyzed the gene expression to better understand the microenvironment of *Cgas^-/-^* mice tumors. We detected the upregulation of *Fn1* (Fig 1F), *Vim* (Fig 1G), *Cxcl12* (Fig 1H), *Il4* (Fig 1I) and *Il10* (Fig 1J). These results suggested that cGAS involves progressive tumorigenesis in MC-38 tumor-bearing mice by increasing vascularity and fibrosis with upregulating *Vim*, a sign of EMT (Epithelial-to-Mesenchymal Transition).

**Fig 1.**
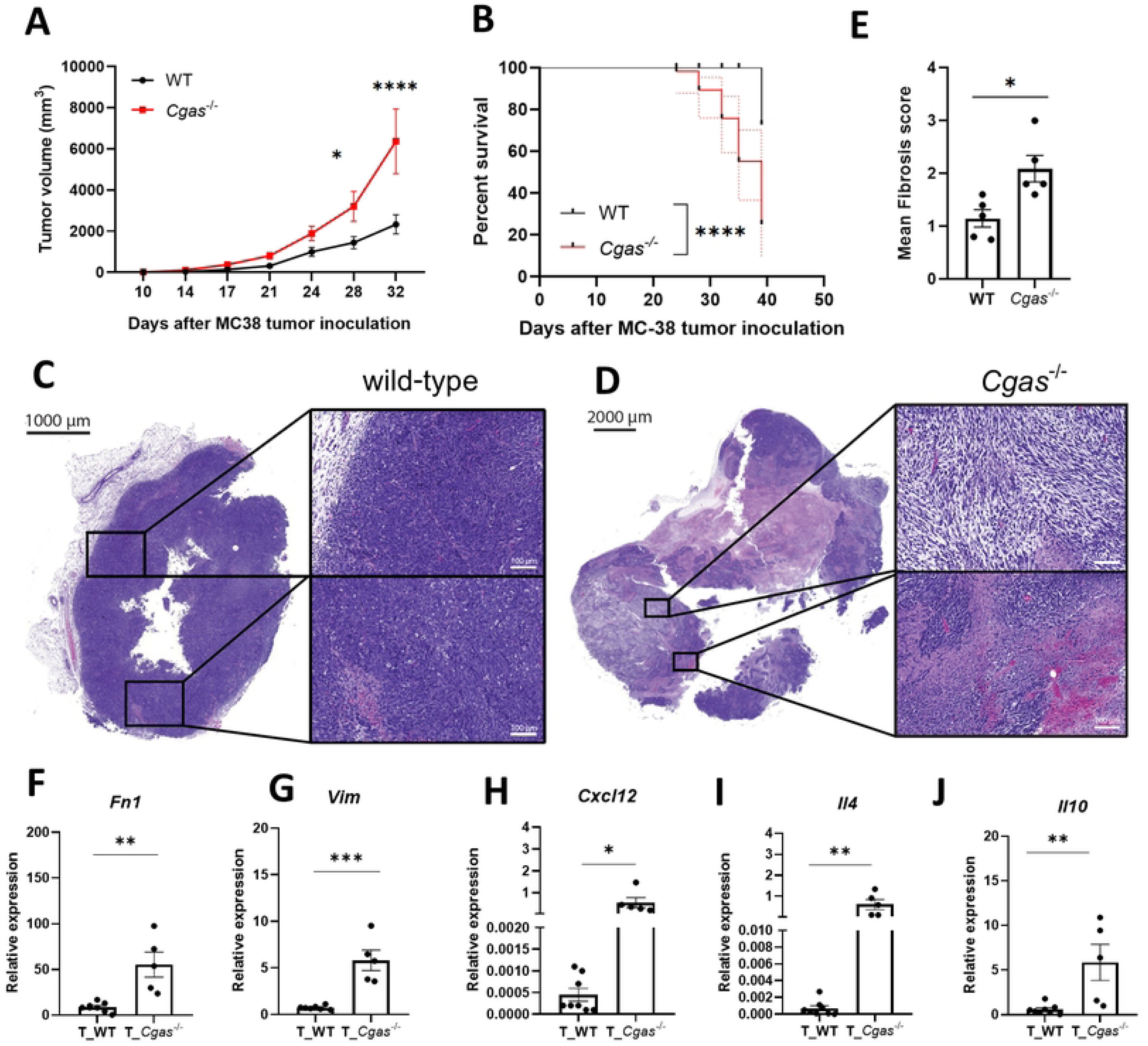
The absence of cGAS promotes progressive tumor formation and poor survival. (A) Mean tumor volume was observed up to 32 days. The statistical data are from two independent experiments, N=12-17 mice per group. (B) The survival rate of mice (N=11 mice per group). (C) Mean fibrosis score of H&E staining. (D-E) FFPE tumor sections of the (D) WT and (E) *Cgas*^-/-^ mice were analyzed by hematoxylin and eosin (H&E) staining (scale bar = 100 mm). Data are representative of 5 mice per group. (F-I) The relative gene expression of (F) *Fn1,* (G) *Vim,* (H) *Cxcl12*, (I) *Il4* and (J) *Il10* genes from the MC-38 tumors of WT and *Cgas*^-/-^ mice. Relative quantities (Δct values) were obtained by normalizing against the *actin* of the spleen control sample. The data are shown as the mean ± SEM (*\*, p < 0.05; **, p < 0.01; ***, p < 0.001*), Student’s *t* test.

### Differential Immune Response in Tumor-Bearing WT and *Cgas^-/-^* Mice

We analyzed the splenocytes to identify the systemic immune response when the mice fully developed tumor progression. We detected the expansion of macrophages and DCs in the spleens of WT and *Cgas*^-/-^ mice (Figs 2A and 2B). However, NK cells in WT mice increased with MC-38 inoculation but not in *Cgas*^-/-^ mice (Fig 2C). At the same time, CD3^+^ and CD4^+^ naïve T cells were reduced in the spleen of both tumor-bearing WT and *Cgas*^-/-^ mice (Figs 2D and 2E), CD4^+^ effector and CD8^+^ effector T cells increased only in tumor-bearing WT mice (Figs 2F and 2G). In contrast, we detected the expansion of CD19^+^ B cells in MC-38-bearing *Cgas*^-/-^ mice (Fig 2H).

**Fig 2.**
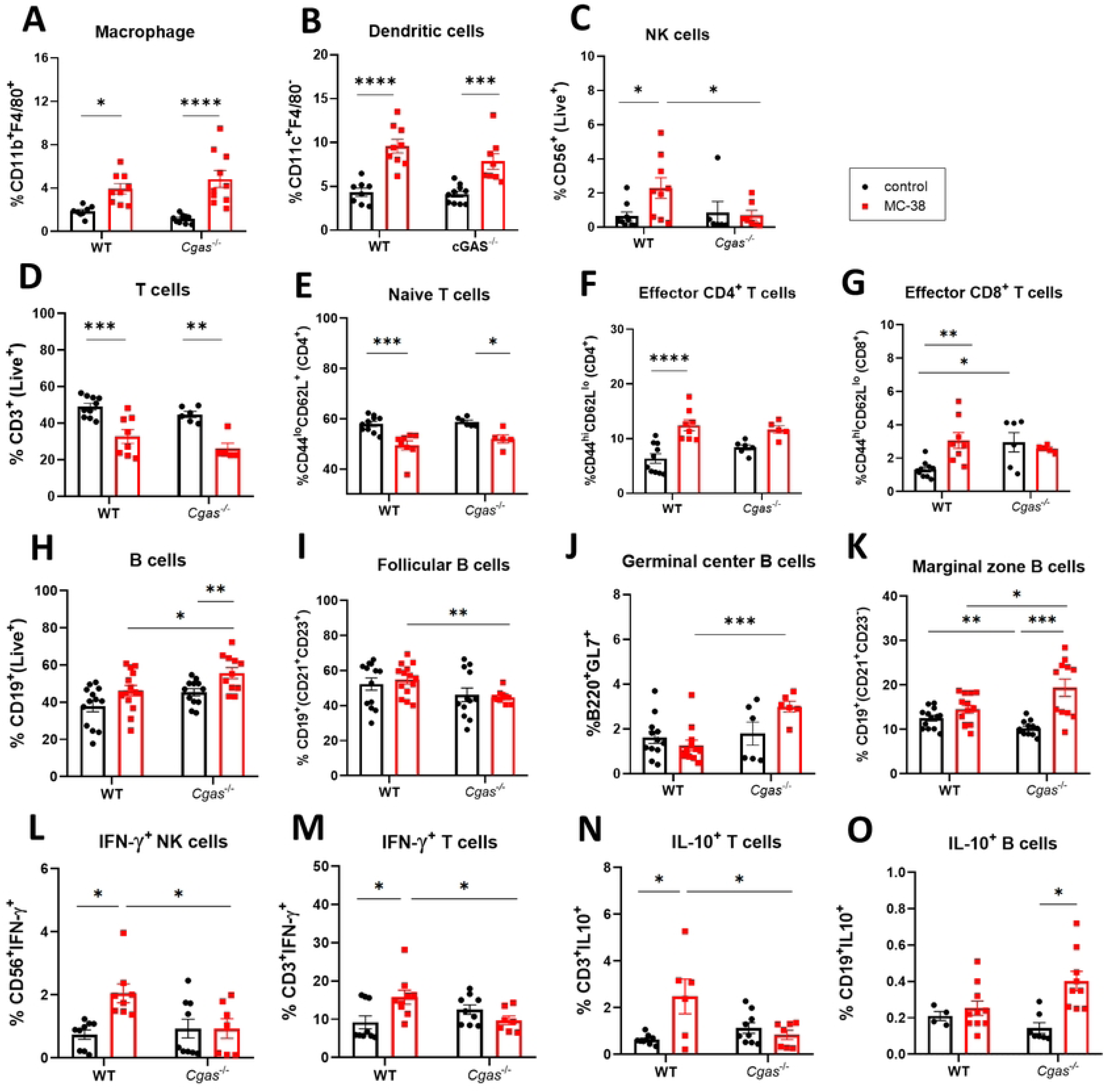
Differential Immune Response in Tumor-Bearing WT and *Cgas^-/-^* Mice. The percentage of (A) macrophages (CD11b^+^F4/80^+^), (B) dendritic cells (CD11c^+^F4/80^-^), (C) natural killer cells (CD56^+^), (D) T cells (CD3^+^), (E) naive T cells (CD3^+^CD4^+^CD44^lo^CD62L^+^), (F) effector CD4^+^ T cells (CD3^+^CD4^+^CD44^hi^CD62L^lo^), (G) effector CD8^+^ T cells (CD3^+^CD8^+^CD44^hi^CD62L^lo^), (H) B cells (CD19^+^), (I) Follicular B cells (CD19^+^CD21^+^CD23^+^), (J) Germinal center B cells (B220^+^GL-7^+^), (K) Marginal zone B cells (CD19^+^CD21^+^CD23^-^), (L) IFN-γ producing-NK cells (CD56^+^IFNγ^+^), (M) IFN-γ producing-T cells (CD3^+^IFNγ^+^), (N) IL-10 producing T cells (CD3^+^IL10^+^) and (O) IL-10 producing B cells (CD19^+^IL10^+^), which were analyzed by flow cytometry from the spleen of the tumor-bearing WT and *Cgas*^-/-^ mice (MC-38) compared to the non-tumor control group (control). N = 5-14 mice per group with 2-3 independent experiments. The statistical data are shown as the mean ± SEM (*, p < 0.05; **, p < 0.01; ***, p < 0.001; ****, p < 0.0001).

There was a reduction in follicular B cells (Fig 2I) but an increase in germinal center B cells (Fig 2J) and marginal zone B cells (Fig 2K) in tumor-bearing *Cgas*^-/-^ mice. Next, we found an induction in IFNγ-producing NK and T cells in tumor-bearing WT mice but not in *Cgas*^-/-^ mice (Figs 2L and 2M). Interestingly, IL-10-producing T cells were increased in tumor-bearing WT mice (Fig 2N), but IL-10-producing B cells (Fig 2O) were increased in tumor-bearing *Cgas*^-/-^ mice. These results suggested that WT mice increased T cell effector function against tumors; in contrast, *Cgas^-/-^* mice showed a response to tumors toward the expansion of B cell subsets and regulatory B cells.

### *cGAS* Deficiency Promotes B Cell Survival in Tumor-Bearing Mice

The expansion of B cells and germinal center B cells were detected in tumor- bearing *Cgas*^-/-^ mice (Figs 2H and 2J). Also, *Cgas*^-/-^ mice exhibited increased autoantibody production in pristane-induced lupus, suggesting that cGAS may regulate hyperactivity of B cell function (19). Thus, we tested the B cell function in tumor- bearing mice. First, we assessed the anti-nuclear antibody (ANA) representing the autoreactivity. We detected higher ANA fluorescent intensity (Figs 3A and 3B) and total IgG levels (Fig 3C) in *Cgas^-/-^* mice without the tumors. Interestingly, MC-38- bearing *Cgas^-/-^* mice showed lower IgG production than WT (Fig 3C). Next, we tested the response of B cells derived from tumor-bearing mice to different stimuli. B cells were cultured with G3-YSD (cGAS agonist) and MC38 TCM, and proliferation was measured using a CFSE dilution assay. The B cells derived from WT and *Cgas*^-/-^ mice showed similar proliferation after being cultured with G3YSD (Figs 3D and 3F) and MC-38 TCM (Figs 3E and 3G). Next, we analyzed the survival of B cells in the culture. The CD19^+^ B cells in tumor-bearing mice survived better in WT and *Cgas*^-/-^ mice than B cells from normal control mice, independent of G3-YSD stimulation (Fig 3H). Then, we cultured B cells with an MC-38 TCM and observed that the survival of *Cgas*^-/-^ B cells derived from tumor-bearing mice was higher than that of WT B cells. In addition, MC-38 TCM boosted the survival of *Cgas*^-/-^ B cells (Fig 3I). These data suggested that the MC-38-bearing environment enhanced intrinsic B cell survival in *Cgas*^-/-^ mice.

**Fig 3.**
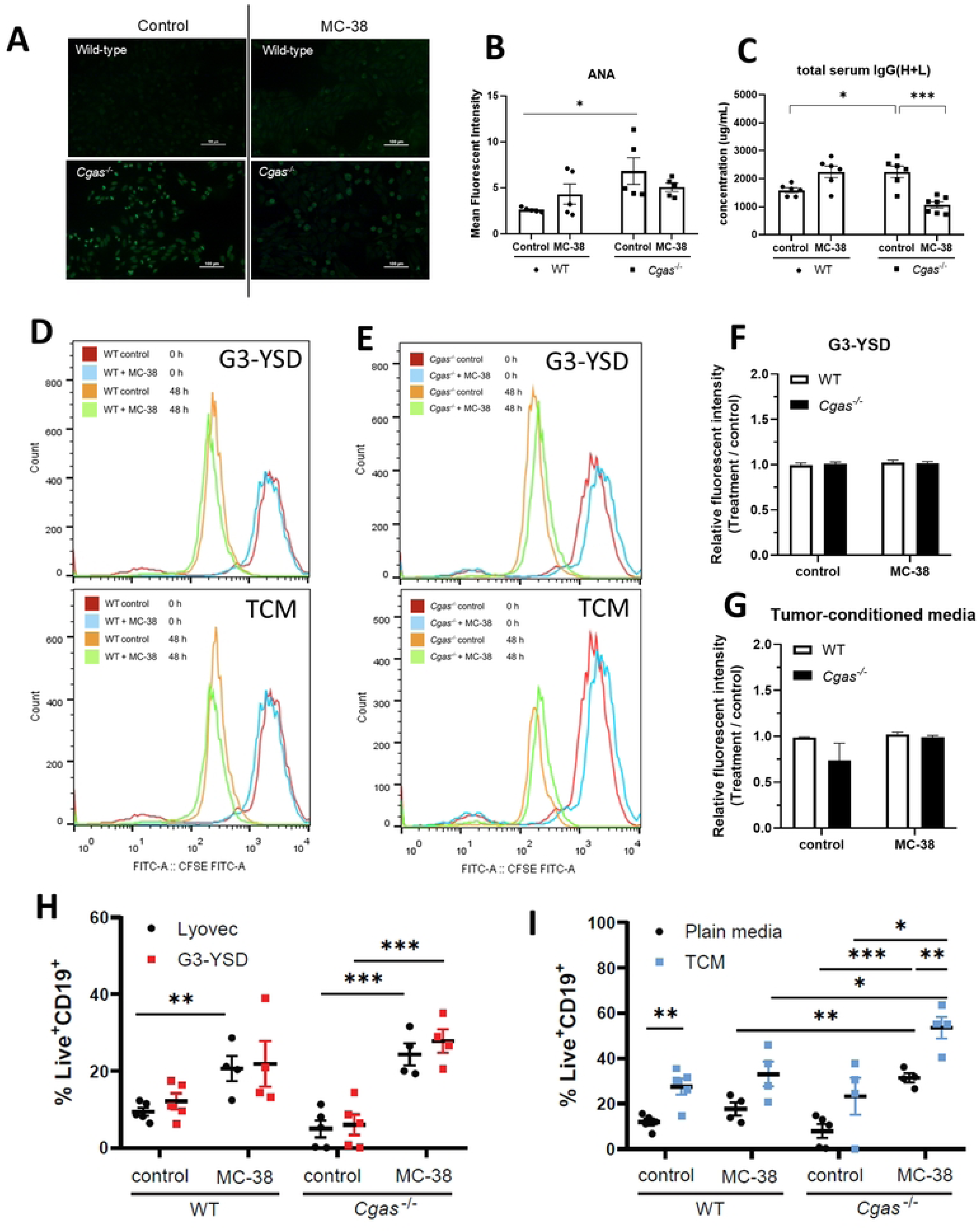
*cGAS* Deficiency Promotes B Cell Survival in Tumor-Bearing Mice. (A) ANAs from mouse sera at a 1:50 dilution. Representative data from 3 mice per group are shown at 200X magnification (scale bar, 100 μm). (B) Samples were analyzed using the mean density of (A). (C) Total serum IgG levels in WT and *Cgas*^-/-^ mice. (D-E) The histogram of CFSE-labeled B cells after treatment with G3-YSD and TCM for 48 hours. (F-G) Mean fluorescent intensity of CFSE. (H-I), The percentage of living B cells after treatment with G3-YSD (H) and tumor-conditioned media (I) from WT mice and *Cgas*^-/-^ mice with/without tumors. The statistical analysis is shown as the mean ± SEM (N= 4-5 per group, 4 independent experiments; *, p < 0.05; **, p < 0.01; ***, p < 0.001).

### cGAS Deficiency Alters Tumor Response to B Cell Depletion by Promoting a Fibrotic and Angiogenic Tumor Microenvironment

B cells can support tumor growth or contribute to antitumor responses (26).

Increased IL-10-producing B cells in tumor-bearing *Cgas^-/-^* mice indicated that regulatory B cells might facilitate tumor progression. To test this, WT and *Cgas^-/-^* mice received an anti-CD20 mAb intravenously three days before tumor implantation. B cell depletion was confirmed in peripheral blood by day 7 post-treatment (S1 Fig). The tumors of B cell-depleted WT mice significantly reduced compared to non-depleted WT mice (Fig 4A). However, B cell-depleted *Cgas^-/-^* mice showed only a trend of tumor volume reduction (Fig 4A). An analysis of MC-38 tumor tissues showed higher fibrosis in B cell-depleted *Cgas^-/-^* than WT mice (Figs 4B and 4C), indicating fibrous tissue as a critical component of residual tumors in *Cgas^-/-^* mice post-depletion.

**Fig 4.**
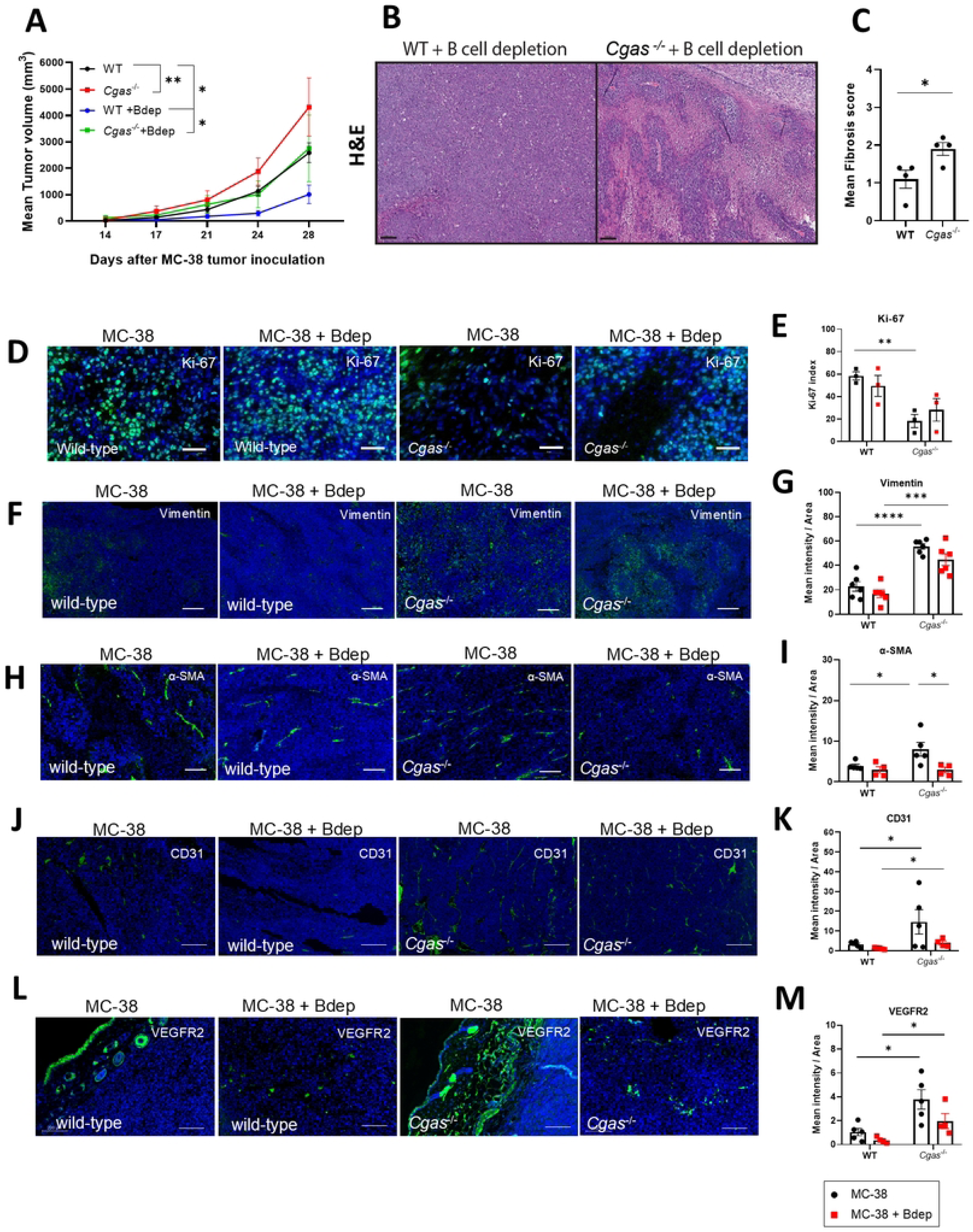
cGAS Deficiency Alters Tumor Response to B Cell Depletion by Promoting a Fibrotic and Angiogenic Tumor Microenvironment. (A) The tumor volume was observed for up to 28 days (N = 5-8 mice per group). (B) MC-38 tumors from B cell-depleted WT and *Cgas^-/-^* mice were stained with H&E. Data represent 4 mice per group. (C) Mean fibrosis score of (B). (D) Immunofluorescence of Ki-67 staining in tumors. Representative data from 3 mice per group are shown at 200x magnification (scale bar=50 μm). (E) Ki-67 index of (D) (N=3 mice per group). (F) Immunofluorescence of Vimentin^+^ tumors. (G) Mean intensity of Vimentin^+^ tumors. (H) Immunofluorescence of α-SMA^+^ tumors.

Next, we examined the characteristics of tumors after B cell depletion. B cell depletion did not affect the Ki-67 expression in tumors of WT and *Cgas^-/-^* mice (Figs 4D and 4E). However, the Ki-67 staining was higher in WT than in *Cgas^-/-^* mice in the non-depleted group. Next, we identified the Epithelial-to-Mesenchymal Transition (EMT) by staining vimentin. We detected higher fluorescence intensity of vimentin in *Cgas^-/-^* mice tumors than WT, which did not change after B cell depletion (Figs 4F and 4G). α-SMA staining revealed increased expression in *Cgas^-/-^* mice’s tumors, significantly reduced by B cell depletion (Figs 4H and 4I). We analyzed CD31 (PECAM-1), a marker of vascular endothelial cells (29), to assess the microvessel density (MVD) in the tumor tissue. *Cgas^-/-^* mouse tumors showed CD31 staining more than WT mice but did not significantly reduce after B cell depletion (Figs 4J and 4K). We also detected increased VEGFR2 staining in *Cgas^-/-^* mouse blood vessels (Figs 4L and 4M). These findings suggest that *Cgas^-/-^* mice contribute to tumor progression through mechanisms involving fibrosis, angiogenesis, and EMT. However, B cell depletion affects tumor stroma and vasculature differently in WT and *Cgas^-/-^* mice.

Representative data from 4-5 mice per group are shown at 100x magnification (scale bar, 200 μm). (I) Mean intensity of α-SMA^+^ tumors. (J) Immunofluorescence of CD31^+^ tumors. (K) Mean intensity of CD31^+^ tumors. (L) Immunofluorescence of VEGFR2^+^ tumors. (M) Mean intensity of VEGFR2^+^ tumors. Representative data from 4-5 mice per group is shown at 100x magnification (scale bar, 200 μm). The statistical analysis is the mean ± SEM (*, p < 0.05; **, p < 0.01).

### cGAS Deficiency Enhances B Cell Survival Mechanisms and Limits CD8^+^ T Cell Infiltration in Tumors

Tumor-infiltrating lymphocytes (TIL) are crucial for controlling tumor growth.

Here, we identified CD8^+^ cells increased in the tumors of WT mice with B cell depletion but not in *Cgas^-/-^* mice. However, the CD8^+^ T cells in MC-38 tumors significantly increased in *Cgas^-/-^* mice compared to WT mice (Figs 5A and 5B). Despite B cell depletion, B cells persisted in tumors of *Cgas^-/-^* mice, while WT showed a notable decrease (Figs 5C and 5D). Next, we tested the factors that make the B cells more resistant to B cell depletion by looking at the signaling and factors for B cell survival. Our study revealed increased expression of *Tgfb1* and *Cd40l* in tumors derived from *Cgas^-/-^* mice, which decreased after B-cell depletion (Figs 5E and 5F).

**Fig 5.**
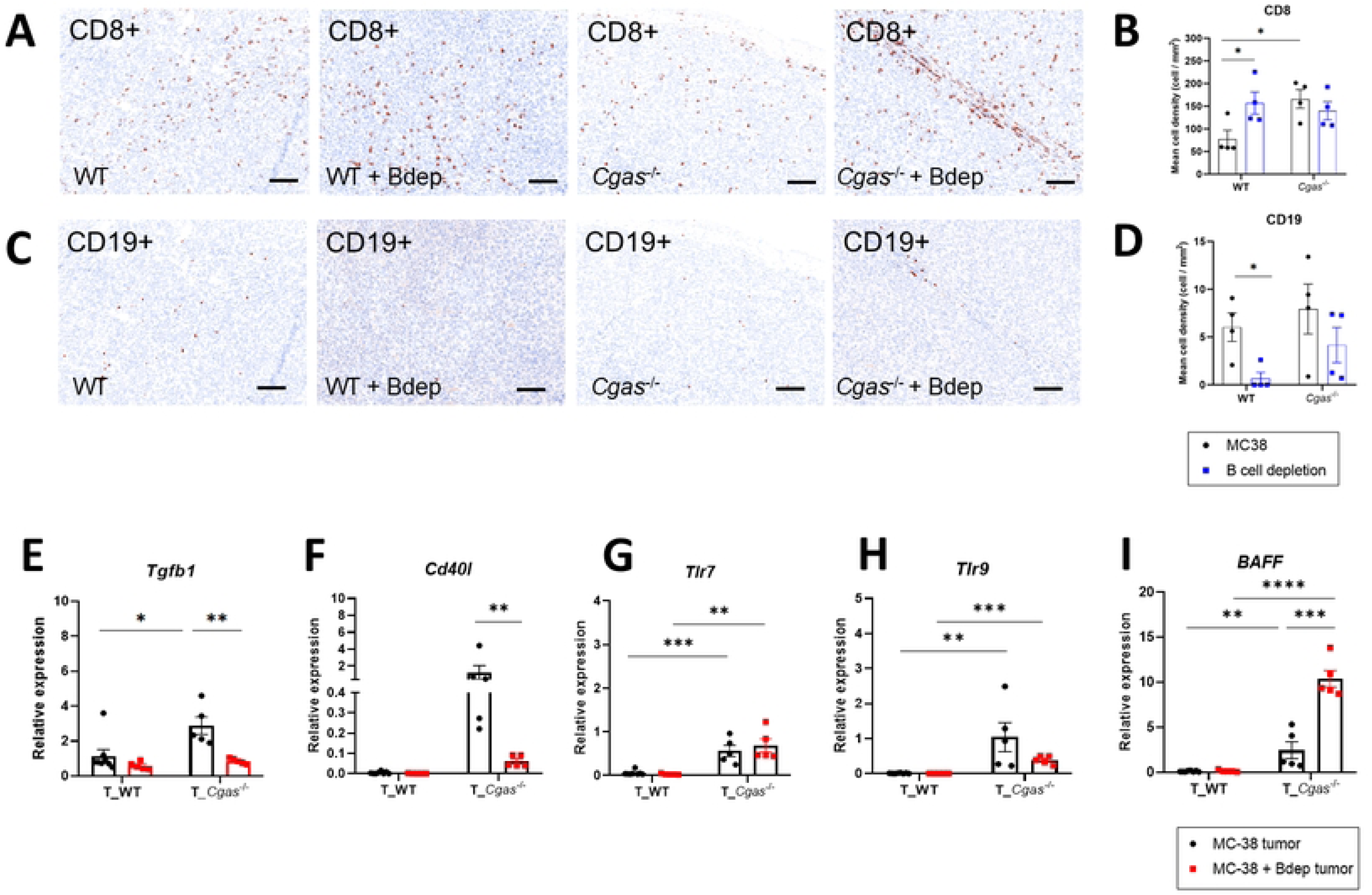
cGAS Deficiency Enhances B Cell Survival Mechanisms and Limits CD8^+^ T Cell Infiltration in Tumors. Immunoprofiling of tumor tissues was analyzed by multiplex immunofluorescence (mIF) (scale bar=100 μm). (A) Tumor staining of CD8^+^ cells. (B) The mean cell densities of CD8^+^ cells. (C) Tumor staining of CD19^+^ cells. (D) The mean cell densities of CD19^+^ cells. The samples were harvested from the tumors of the B cell- depleted mice compared to the non-depleted mice (N = 3-4 mice per group). The density of the staining cell was quantified by inForm® Tissue Analysis Software. The bar is shown as the mean ± SEM (*, p < 0.05; **, p < 0.01; ***, p < 0.001; ****, p < 0.0001).

Expression of *Tlr7* and *Tlr9* was higher in *Cgas^-/-^* mice and was unaffected by B cell depletion (Figs 5G and 5H). Interestingly, *Baff* expression was higher in *Cgas^-/-^* mouse tumors and increased after B cell depletion (Fig 5I). The data suggest that cGAS deficiency promotes a tumor microenvironment that supports B cell survival and resistance to depletion, potentially through increased expression of survival factors such as BAFF and persistent signaling via Tlr7 and Tlr9.

### cGAS Deficiency in B Cells Enhances Blood Vessel Formation

Since we detected more blood vessels in *Cgas^-/-^* mice tumors and B cell depletion reduced the expression of α-SMA, we hypothesized that B cells from *Cgas^-/-^* mice may promote angiogenesis, which could enhance tumor progression. To identify whether cGAS-deficient B cells are positively involved in angiogenesis, we co-cultured isolated B cells from MC-38-bearing WT and *Cgas^-/-^* mice with HUVECs to study endothelial tube formation assay. The result revealed that *Cgas^-/-^* B cells significantly induced more endothelial tube formation than WT B cells (Fig 6A), considerably increasing in nodes and branches at 4 hours (Figs 6B and 6C). These findings suggest that *Cgas^-/-^* B cells could contribute to tumor progression by enhancing blood vessel formation, potentially supporting increased tumor growth.

**Fig 6.**
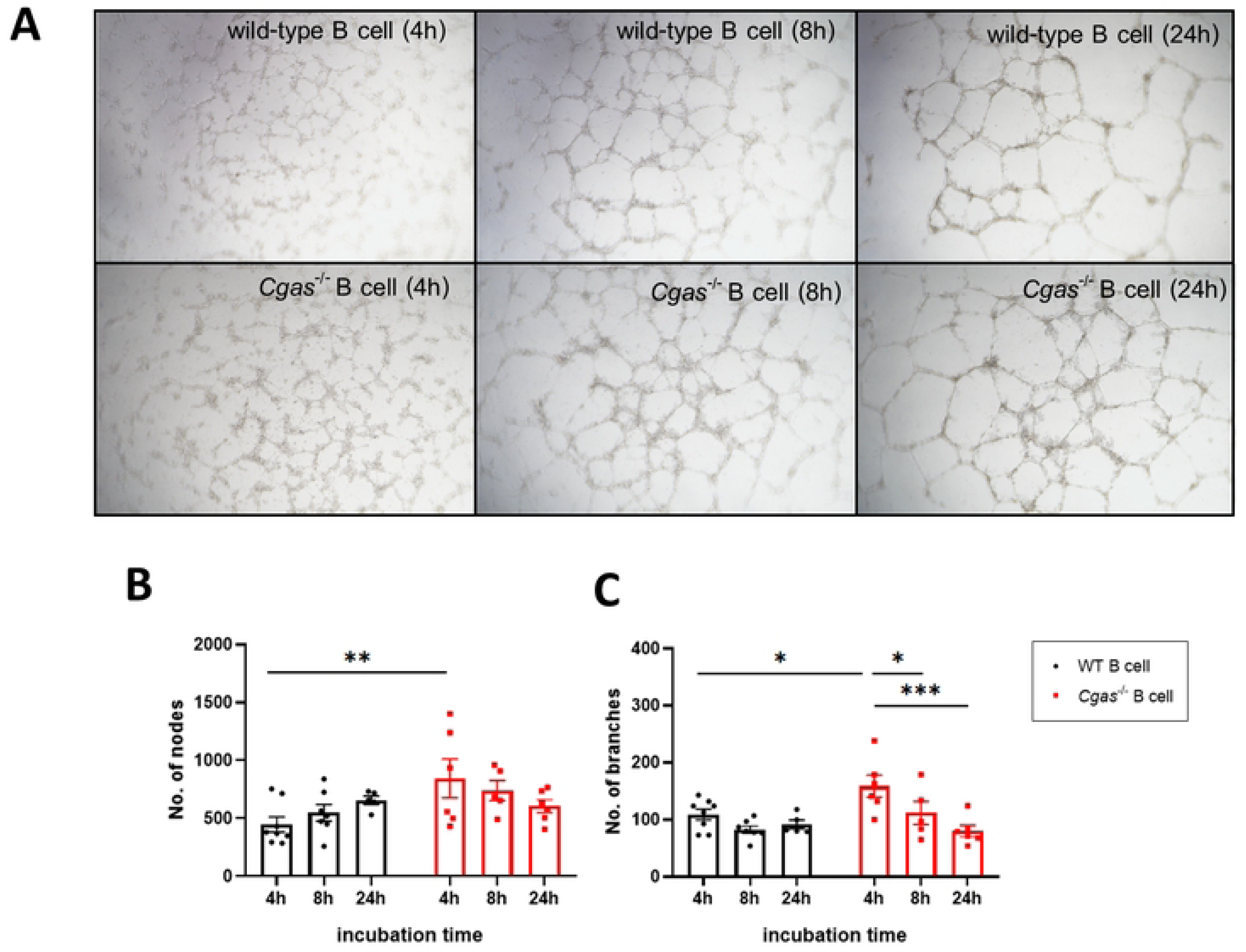
cGAS Deficiency in B Cells Enhances Blood Vessel Formation. B cells derived from MC-38-bearing WT and *Cgas^-/-^* mice were cultured with HUVECS up to 24 hours. (A) Figures showed tube formation in Matrigel-coated wells by HUVECs cocultured with B cells. Representative data from 6-8 mice per group (2 independent experiments) are shown (scale bar= 200 μm). (B) The number of nodes and (C) the number of branches were analyzed by ImageJ. The bar is shown as the mean ± SEM (*, p < 0.05; **, p < 0.01; ***, p < 0.001).

## Discussion

cGAS senses tumor DNA, and its absence facilitates immune escape and tumorigenesis (30). Lack of the *Cgas* gene was linked to cancer in a mouse model of AOM-induced colon cancer (31). Our study found cGAS signaling essential for slowing MC-38 tumor growth and extending survival. B cells in *Cgas^-/-^* mice might promote angiogenesis by upregulating proangiogenic cytokines like VEGF, FGF2, and PDGFA (32). In addition, the histology of tumor cells did not increase proliferation or the Ki-67 index in *Cgas*^-/-^ mice, but fibrosis, vascularity, and EMT were dramatically present. Advanced tumor growth in *Cgas*^-/-^ mice could occur from the TME-promoting fibrosis, angiogenesis, and EMT, leading to aggressive phenotypes. Additionally, the increase of Il10 and Il4 expression could suggest the immunosuppressive tumor microenvironment (TME) of *Cgas^-/-^* mice, which could enhance tumor progression.

Immune responses changed in tumor-bearing mice. Activating effector T cells and IFN-γ production attack tumor antigens and mediate cytotoxic reactions (6, 33). WT mice showed increased NK cells, effector T cells, and IFN-γ, while *Cgas^-/-^* mice had more marginal zone B cells and IL-10-producing B cells, indicating a B regulatory cell (Breg) phenotype. Marginal zone B cells in *Cgas^-/-^* mice produced IL-10, suppressing T cell effector function against tumors. Bregs can inhibit antitumor T cells and promote tumor growth (34).

Antinuclear antibodies (ANAs) detected in cancers can influence treatment prognosis (35). Increased plasma cells and germinal center B cells in tumor-bearing *Cgas^-/-^* mice indicate tumor antigen-driven B cell reactivity and ANA detection. B cells can produce autoantibodies via intrinsic or extrinsic activating factors (36). B cells from *Cgas^-/-^* mice showed enhanced survival in tumor-medium conditions, highlighting the tumor microenvironment’s (TME) role in promoting B cell survival. Increased *Baff* expression in tumors points to BAFF-producing cell infiltration in tumors. BAFF stimulates B cell differentiation, survival, and Ig production and may originate from tumor-associated macrophages (37, 38), suggesting BAFF from these macrophages enhances B cell survival and tumor growth in *Cgas^-/-^* mice.

Patients with autoimmune diseases are at higher risk of cancer (39). Circulating immune complexes (CICs) indicate poor cancer prognosis (40). B cell depletion therapy is preferred for B cell malignancies. However, B cell depletion in solid tumors may slow progression without inducing remission (41). The limited success of B cell depletion in cancer treatment might stem from the persistence of chronic stimulation with immunoglobulins or CICs, which do not vanish immediately after B cell depletion.

A study found that B cell depletion on day 9 post-tumor injection did not reduce MC-38 tumor volume (42). However, depleting B cells 3 days before inoculation significantly lowered MC-38 tumor volume, highlighting B cells’ role in early tumor development. Higher ANA and B cell levels in *Cgas^-/-^* mice with MC-38 tumors than in WT mice suggest that *Cgas^-/-^* mice’s autoreactivity may promote tumor growth. B cell depletion partially reduced tumor progression in *Cgas^-/-^* mice. MC-38 tumors’ extensive fibrosis in *Cgas^-/-^* mice might hinder anti-CD20 mAb delivery. Also, B cells in *Cgas^-/-^* mice survived longer, causing resistance to B cell depletion in these mice.

Fibrosis in the TME helps tumors evade immune defense. TGF-β is crucial in tumorigenesis, promoting EMT, cell proliferation, invasion, metastasis, angiogenesis, and immune evasion (43). Reducing TGF-β in *Cgas^-/-^* mice tumors after B cell depletion indicates a significant role of B cells in TGF-β production. TGF-β contributes to tumorigenesis, including tumor growth, fibrogenesis, and angiogenesis. (44).

Regulatory B cells secrete TGF-β to control inflammation in experimental autoimmune encephalomyelitis (EAE) (45). Moreover, TGF-β^+^ Bregs and IL-10 are found in various tumor environments, contributing to tumor growth and metastasis (46).

However, B cell depletion did not reduce IL-10 in MC-38 tumors in *Cgas^-/-^* mice, suggesting T regulatory cells might also produce IL-10 in tumors.

Inflammatory cytokines influence cancer treatment outcomes. Among 125 NSCLC patients, those with low IL-6 had better disease control with immunotherapy than those with high IL-6 (47). Increasing *Il1a*, *Il6*, and *Tnfa* expression in MC-38 tumors in *Cgas*^-/-^ mice suggested a proinflammatory TME, which could enhance tumor development and progression, metastasis, and resistance to chemotherapy. The presence of *Tlr7*, *Tlr9*, and *Baff* in *Cgas*^-/-^ tumors gave survival factors to B cells and may make B cells more resistant to being depleted. The persistence of these cytokines after B cell depletion in *Cgas^-/-^* mice implies that other cell types also produce these cytokines. B cell depletion reducing new blood vessel formation in MC-38 TME of *Cgas****^-^****^/^*^-^ mice shows B cells’ role in angiogenesis. Proangiogenic B cells increase the expression of *TGFB2* and *VEGFA* (32). Reduced *Tgfb* and *Vegfa* in splenocytes post-B cell depletion highlights the proangiogenic nature of B cells from MC-38-bearing *Cgas^-/-^* mice.

## Conclusions

In summary, *Cgas*^-/-^ mice exhibit progressive tumor growth characterized by angiogenesis, fibrosis, increased regulatory B cells (Bregs), EMT, and an elevated proinflammatory tumor microenvironment (TME). The expansion of Bregs and proangiogenic B cells in *Cgas*^-/-^ mice likely impedes the natural immune response against cancer. Additionally, the development of autoantibodies in tumor-bearing mice suggests that B cell autoreactivity occurs, and detecting autoantibodies in cancer patients may serve as a potential biomarker for B cell depletion as an adjunct immunotherapy. cGAS deficiency fosters a microenvironment where B cells not only survive but also actively contribute to tumor angiogenesis, further promoting tumor growth and progression. The dual role of B cells in immune suppression and vascular support positions them as critical players in cGAS-deficient tumorigenesis, making them a potential therapeutic target in solid cancers. Strategies to deplete B cells or block their survival pathways, such as BAFF or TLR signaling, could disrupt the immunosuppressive and angiogenic functions supporting tumor growth. Further research is needed to identify patients with specific biomarkers, such as ANA, to determine appropriate candidates for B cell depletion therapy and to explore combination therapies to overcome resistance to B cell depletion observed in cGAS- deficient mice.

## Acknowledgments

We want to thank Professor Søren Riis Paludan for providing the *Cgas*-deficient mice, all CUSB staff, and the Faculty of Medicine, Chulalongkorn University animal center team for help and support.

## Supporting Information

S1 Table. qRT-PCR primers

S1 Fig. The percentage of B cells in peripheral blood at day 7 after B cell depletion.

## Data availability Statement

All data generated or analyzed during this study are included in this published article [and its supplementary information files].

## Funding

This research was funded by the Science Achievement Scholarship of Thailand (SAST) granted to PC, Fundamental Fund, FRB65_hea(44)_051_30_32., the Research University Network (RUN), PMI06/2561 from the National Research Council of Thailand (NRCT), Thailand granted to TP.

## Author Contributions

Conceptualization: Papasara Chantawichitwong, Prapaporn Pisitkun, Trairak Pisitkun

Data curation: Papasara Chantawichitwong

Formal analysis: Papasara Chantawichitwong, Prapaporn Pisitkun

Funding acquisition: Prapaporn Pisitkun, Trairak Pisitkun, Papasara Chantawichitwong

Investigation: Papasara Chantawichitwong, Sarinya Kumpunya, Tossapon Wongtangprasert, Peerapat Visitchanakun, Prapaporn Pisitkun

Methodology: Papasara Chantawichitwong, Prapaporn Pisitkun, Trairak Pisitkun

Project administration: Prapaporn Pisitkun, Trairak Pisitkun

Resources: Prapaporn Pisitkun, Trairak Pisitkun

Software: Papasara Chantawichitwong, Prapaporn Pisitkun Supervision: Prapaporn Pisitkun, Trairak Pisitkun Validation: Papasara Chantawichitwong, Prapaporn Pisitkun Visualization: Papasara Chantawichitwong

Writing – original draft: Papasara Chantawichitwong

Writing – review & editing: Papasara Chantawichitwong, Prapaporn Pisitkun, Trairak Pisitkun

PP and TP are joint senior authors

## Declaration of competing interest

The authors declare that there are no conflicts of interest.

## Notes

### Competing Interest Statement

The authors have declared no competing interest.

## References

1. Whiteside TL. The tumor microenvironment and its role in promoting tumor growth. Oncogene. 2008;27(45):5904–12.

2. Jeong H, Hwang I, Kang SH, Shin HC, and Kwon SY. Tumor-Associated Macrophages as Potential Prognostic Biomarkers of Invasive Breast Cancer. J Breast Cancer. 2019;22(1):38–51.

3. Gao G, Wang Z, Qu X, and Zhang Z. Prognostic value of tumor-infiltrating lymphocytes in patients with triple-negative breast cancer: a systematic review and meta-analysis. BMC Cancer. 2020;20(1):179.

4. Idos GE, Kwok J, Bonthala N, Kysh L, Gruber SB, and Qu C. The Prognostic Implications of Tumor Infiltrating Lymphocytes in Colorectal Cancer: A Systematic Review and Meta-Analysis. Sci Rep. 2020;10(1):3360.

5. Michaud D, Steward CR, Mirlekar B, and Pylayeva-Gupta Y. Regulatory B cells in cancer. Immunol Rev. 2021;299(1):74–92.

6. Sun L, Wu J, Du F, Chen X, and Chen ZJ. Cyclic GMP-AMP synthase is a cytosolic DNA sensor that activates the type I interferon pathway. Science. 2013;339(6121):786-91.

7. Li T, and Chen ZJ. The cGAS-cGAMP-STING pathway connects DNA damage to inflammation, senescence, and cancer. J Exp Med. 2018;215(5):1287–99.

8. Vanpouille-Box C, Alard A, Aryankalayil MJ, Sarfraz Y, Diamond JM, Schneider RJ, et al. DNA exonuclease Trex1 regulates radiotherapy-induced tumour immunogenicity. Nat Commun. 2017;8:15618.

9. Shen R, Liu D, Wang X, Guo Z, Sun H, Song Y, et al. DNA Damage and Activation of cGAS/STING Pathway Induce Tumor Microenvironment Remodeling. Front Cell Dev Biol. 2021;9:828657.

10. Li J, and Bakhoum SF. The pleiotropic roles of cGAS-STING signaling in the tumor microenvironment. J Mol Cell Biol. 2022;14(4).

11. Zhao H, Wu L, Yan G, Chen Y, Zhou M, Wu Y, et al. Inflammation and tumor progression: signaling pathways and targeted intervention. Signal Transduct Target Ther. 2021;6(1):263.

12. Elinav E, Nowarski R, Thaiss CA, Hu B, Jin C, and Flavell RA. Inflammation-induced cancer: crosstalk between tumours, immune cells and microorganisms. Nat Rev Cancer. 2013;13(11):759–71.

13. Sauleda J, Nunez B, Sala E, and Soriano JB. Idiopathic Pulmonary Fibrosis: Epidemiology, Natural History, Phenotypes. Med Sci (Basel). 2018;6(4).

14. Wu B, Sodji QH, and Oyelere AK. Inflammation, Fibrosis and Cancer: Mechanisms, Therapeutic Options and Challenges. Cancers (Basel). 2022;14(3).

15. Xu J, Lamouille S, and Derynck R. TGF-beta-induced epithelial to mesenchymal transition. Cell Res. 2009;19(2):156–72.

16. Ota T, Fujii M, Sugizaki T, Ishii M, Miyazawa K, Aburatani H, et al. Targets of transcriptional regulation by two distinct type I receptors for transforming growth factor-beta in human umbilical vein endothelial cells. J Cell Physiol. 2002;193(3):299–318.

17. Shi X, Yang J, Deng S, Xu H, Wu D, Zeng Q, et al. TGF-beta signaling in the tumor metabolic microenvironment and targeted therapies. J Hematol Oncol. 2022;15(1):135.

18. Kitai Y, Kawasaki T, Sueyoshi T, Kobiyama K, Ishii KJ, Zou J, et al. DNA-Containing Exosomes Derived from Cancer Cells Treated with Topotecan Activate a STING-Dependent Pathway and Reinforce Antitumor Immunity. J Immunol. 2017;198(4):1649–59.

19. Liu X, Pu Y, Cron K, Deng L, Kline J, Frazier WA, et al. CD47 blockade triggers T cell-mediated destruction of immunogenic tumors. Nat Med. 2015;21(10):1209–15.

20. Li S, Mirlekar B, Johnson BM, Brickey WJ, Wrobel JA, Yang N, et al. STING-induced regulatory B cells compromise NK function in cancer immunity. Nature. 2022;610(7931):373-80.

21. Abu-Shakra M, Buskila D, Ehrenfeld M, Conrad K, and Shoenfeld Y. Cancer and autoimmunity: autoimmune and rheumatic features in patients with malignancies. Ann Rheum Dis. 2001;60(5):433–41.

22. Thim-Uam A, Prabakaran T, Tansakul M, Makjaroen J, Wongkongkathep P, Chantaravisoot N, et al. STING Mediates Lupus via the Activation of Conventional Dendritic Cell Maturation and Plasmacytoid Dendritic Cell Differentiation. iScience. 2020;23(9):101530.

23. Kumpunya S, Thim-Uam A, Thumarat C, Leelahavanichkul A, Kalpongnukul N, Chantaravisoot N, et al. cGAS deficiency enhances inflammasome activation in macrophages and inflammatory pathology in pristane-induced lupus. Front Immunol. 2022;13:1010764.

24. Sakimura C, Tanaka H, Okuno T, Hiramatsu S, Muguruma K, Hirakawa K, et al. B cells in tertiary lymphoid structures are associated with favorable prognosis in gastric cancer. J Surg Res. 2017;215:74–82.

25. Bindea G, Mlecnik B, Tosolini M, Kirilovsky A, Waldner M, Obenauf AC, et al. Spatiotemporal dynamics of intratumoral immune cells reveal the immune landscape in human cancer. Immunity. 2013;39(4):782–95.

26. Wang SS, Liu W, Ly D, Xu H, Qu L, and Zhang L. Tumor-infiltrating B cells: their role and application in anti-tumor immunity in lung cancer. Cell Mol Immunol. 2019;16(1):6–18.

27. Andreu P, Johansson M, Affara NI, Pucci F, Tan T, Junankar S, et al. FcRgamma activation regulates inflammation-associated squamous carcinogenesis. Cancer Cell. 2010;17(2):121–34.

28. Motwani M, McGowan J, Antonovitch J, Gao KM, Jiang Z, Sharma S, et al. cGAS-STING Pathway Does Not Promote Autoimmunity in Murine Models of SLE. Front Immunol. 2021;12:605930.

29. Guo CR, Han R, Xue F, Xu L, Ren WG, Li M, et al. Expression and clinical significance of CD31, CD34, and CD105 in pulmonary ground glass nodules with different vascular manifestations on CT. Front Oncol. 2022;12:956451.

30. Schadt L, Sparano C, Schweiger NA, Silina K, Cecconi V, Lucchiari G, et al. Cancer-Cell-Intrinsic cGAS Expression Mediates Tumor Immunogenicity. Cell Rep. 2019;29(5):1236–48 e7.

31. Hu S, Fang Y, Chen X, Cheng T, Zhao M, Du M, et al. cGAS restricts colon cancer development by protecting intestinal barrier integrity. Proc Natl Acad Sci U S A. 2021;118(23).

32. van de Veen W, Globinska A, Jansen K, Straumann A, Kubo T, Verschoor D, et al. A novel proangiogenic B cell subset is increased in cancer and chronic inflammation. Sci Adv. 2020;6(20):eaaz3559.

33. Mizuno R, Kawada K, Itatani Y, Ogawa R, Kiyasu Y, and Sakai Y. The Role of Tumor-Associated Neutrophils in Colorectal Cancer. Int J Mol Sci. 2019;20(3).

34. Catalán D, Mansilla MA, Ferrier A, Soto L, Oleinika K, Aguillón JC, et al. Immunosuppressive Mechanisms of Regulatory B Cells. Front Immunol. 2021;12:611795.

35. Vlagea A, Falagan S, Gutiérrez-Gutiérrez G, Moreno-Rubio J, Merino M, Zambrana F, et al. Antinuclear antibodies and cancer: A literature review. Crit Rev Oncol Hematol. 2018;127:42–9.

36. Suurmond J, and Diamond B. Autoantibodies in systemic autoimmune diseases: specificity and pathogenicity. J Clin Invest. 2015;125(6):2194–202.

37. Sun J, Park C, Guenthner N, Gurley S, Zhang L, Lubben B, et al. Tumor-associated macrophages in multiple myeloma: advances in biology and therapy. J Immunother Cancer. 2022;10(4).

38. Moore PA, Belvedere O, Orr A, Pieri K, LaFleur DW, Feng P, et al. BLyS: member of the tumor necrosis factor family and B lymphocyte stimulator. Science. 1999;285(5425):260-3.

39. Li CM, and Chen Z. Autoimmunity as an Etiological Factor of Cancer: The Transformative Potential of Chronic Type 2 Inflammation. Front Cell Dev Biol. 2021;9:664305.

40. Rai S, and Mody RN. Serum circulating immune complexes as prognostic indicators in premalignant and malignant lesions of oral cavity during and following radiotherapy. J Cancer Res Ther. 2012;8 Suppl 1:S116–22.

41. Kim S, Fridlender ZG, Dunn R, Kehry MR, Kapoor V, Blouin A, et al. B-cell depletion using an anti-CD20 antibody augments antitumor immune responses and immunotherapy in nonhematopoetic murine tumor models. J Immunother. 2008;31(5):446–57.

42. Damsky W, Jilaveanu L, Turner N, Perry C, Zito C, Tomayko M, et al. B cell depletion or absence does not impede anti-tumor activity of PD-1 inhibitors. J Immunother Cancer. 2019;7(1):153.

43. Baba AB, Rah B, Bhat GR, Mushtaq I, Parveen S, Hassan R, et al. Transforming Growth Factor-Beta (TGF-beta) Signaling in Cancer-A Betrayal Within. Front Pharmacol. 2022;13:791272.

44. Itatani Y, Kawada K, and Sakai Y. Transforming Growth Factor-beta Signaling Pathway in Colorectal Cancer and Its Tumor Microenvironment. Int J Mol Sci. 2019;20(23).

45. Bjarnadottir K, Benkhoucha M, Merkler D, Weber MS, Payne NL, Bernard CCA, et al. B cell-derived transforming growth factor-beta1 expression limits the induction phase of autoimmune neuroinflammation. Sci Rep. 2016;6:34594.

46. Huai G, Markmann JF, Deng S, and Rickert CG. TGF-beta-secreting regulatory B cells: unsung players in immune regulation. Clin Transl Immunology. 2021;10(4):e1270.

47. Kang DH, Park CK, Chung C, Oh IJ, Kim YC, Park D, et al. Baseline Serum Interleukin-6 Levels Predict the Response of Patients with Advanced Non-small Cell Lung Cancer to PD-1/PD-L1 Inhibitors. Immune Netw. 2020;20(3):e27.

